# A new method for measuring fish swimming patterns in shallow water and a previously undescribed swimming gait

**DOI:** 10.1101/853853

**Authors:** Jay Willis, Theresa Burt de Perera, Guillaume Poncelet, Adrian Thomas

## Abstract

Hill stream loaches are fish which live their entire lives in close contact with rock. They have elaborate physical adaptations to fast flow, adherence to substrate, and movement in very shallow water. Here we describe a method for observing how they swim in detail. There are many similarly shaped rheophilic fish, insects, and amphibian larvae, which live in fast flowing water, and a method of observing their swimming modes has wide potential application. We measured the deflection of the water surface around a swimming fish by viewing a fixed pattern on the bottom of the tank through the water surface. This is a Schlieren method in which the movement or other physical properties of a medium are derived from the deflection of a pattern viewed through that medium. We used this method to describe a new type of swimming gait which is likely to be common among small rheophiles – pulse swimming mode – in which thrust is produced in a series of discrete impulses. The method of analysis described here is beneficial in that the fish is allowed to swim freely in relatively normal conditions without the use of intrusive equipment such as lasers, dyes, or additives to the water, and the pattern of thrust is viewed directly against the skin of the fish rather than being inferred from the wake pattern behind the fish. The method is also low cost and easily set up.

## Introduction

The standard approach is to describe open water fish swimming modes as on a spectrum between Anguilliform (like an eel) through Carangiform (like a mackerel) to Tunniform (like a tuna); to distinguish between the sinuous swimming gait of eels and stiff tail based motion of tuna (Breder 1926). Sub-divisions on this spectrum for classification of species, such as sub-carangiform etc., seem contrived but potentially helpful (Webb (1994) described around 20 gaits of swimming fish); some fish can exhibit a range of swimming styles dependent on behaviour, especially for instance when accelerating as opposed to cruising, and especially when confronted with altered flows such as structured turbulence (Liao 2007); and practically all fish alter their suite of potential modes as they grow. A more inclusive perspective is to describe swimming gaits as 1) lift induced (using aerofoils, that conventionally in air are said to provide lift, and in water provide thrust – examples are tuna, billfish and whales), 2) based on undulation (like a trout or a mackerel similar to carangiform and anguilliform above), 3) drag based (using fins or feet as paddles or oars, like frogs or waterbirds such as ducks for instance) and 4) jet propelled (like squid or jellyfish). There are other modes not included in either of these schemes based on the use of undulating fins other than the caudal (knifefish (Apteronotus albifrons)) or, in very small animals, cilia (such as ctenophores). In fish at least, these fin based modes are often used in combination, or addition, with other gaits. Knifefish use fin based swimming for fine control over very short distances and will move backward and forward quickly using this mode, but they are also capable of full body undulation for longer distance more sustained swimming in a forward direction or for rapid escapes. These modes are important with respect to the mode of swimming employed by the fish in this study as these fish appear to swim in a way that does not fit easily into any of these categories, but which may be quite common.

Rheophilic fish such as hill stream loaches are common in fast flowing highland rivers on all continents other than Antarctica. In all cases rheophilic fish interact very closely with the substrate even when physically detached; the hydrodynamic analysis of their swimming is thus complicated by constant presence of heterogeneous flow patterns, turbulence and by the physical characteristics of the substrate. These complexities are often further compounded by rapid depth variations, proximity of the air water interface and entrained bubbles. Rheophilic fish live and swim in a messy hydrodynamic environment far from the simplified cases preferred in mathematical or numerical models. However it is important to note that there are comparatively few studies of fish swimming close to, or almost in contact with, substrates (e.g.: Gerstner 1997) The way fish climb or ‘walk’ over wet or partially submerged substrate has been described with a view to linking these movements with the emergences of animals from the aquatic environment to land in the evolutionary context (Flammang et al 2016).

Schlieren methods are based on the way a transparent medium or object deflects light as light passes through it. Schlieren comes from the German word for streaks, and the effect was first noted by Robert Hooke (1665). Schlieren methods have been used extensively in scientific research for visualising patterns in transparent media such as air or water (Settles 2012). The Schlieren methods used in this study are Background orientated Schlieren methods (Meier 2002) in which a deliberately introduced background is viewed through the transparent medium and the changes in the medium are inferred from the deflection of background. Background orientated Schlieren generally has the great benefit of only requiring a camera and an introduced background, whereas other Schlieren methods involving lasers, discharge lamps, or other light sources viewed through the medium, often require many more optical devices (lenses and detectors) which each introduce error, cost and complexity. In the background orientated method the main burden of analysis is transferred to the computer which is used to determine the deflections across the background pattern in the images captured by the camera.

Digital Image Correlation (DIC) was first developed in the 1980’s and has been used extensively in the field of stress research (Hild and Roux 2006). DIC works on the principle of comparison of two photographs taken a short time apart. The process compares groups of pixels on one image (the reference image) with those on the other (the current image) and searches for similarities from which the movement of the pixels from one image to the other is calculated. In stress research it allows very small strains over a wide area of a solid flexible body to be observed in a way that might be difficult with a set of fixed strain gauges. The mathematics of the process show how movements below the spatial scale of a single pixel can be detected and how errors can be reliably estimated (Hild and Roux 2006). The process only requires two photographs as a minimum, but can be extended to a sequence of current images off a baseline reference image which can allow for smoothing of the derived displacement fields, using multiple images to quantify the mean likely displacement field in each image and the associated bias across multiple images. Since the method has been used extensively over many years, the algorithms are well developed, but the process remains extremely computer intensive. The steady improvement of desktop computing power over the past few years has led to DIC analysis only recently becoming something that can be reliably be performed by a non-specialist scientist without custom computer hardware.

In this study we use Background Orientated Schlieren to observe the defections in a background image produced by waves on the surface of water in a shallow tank. In our case the waves have been produced by a swimming fish and the background pattern is visible through the base of the glass tank. The technique of deriving the surface wave pattern from observation of the basal pattern, in a similar set up to the one described here, has been introduced and explained by Moisy et al. (2009).

## Methods

### Apparatus and set up

We used a Sony RX10ii Camera (sony.com) suspended 1 m above a shallow tank of water. The basal pattern was made using software provided by Moisy et al. (2009) using MATLAB (mathworks.com). It was printed on A4 paper and laminated. The tank was custom built from 6 mm float glass and silicon sealant; it was 0.65 m x 0.30 m x 0.15 m in external dimension, manufactured by Clear Seal (clearseal.com). The water depth was varied between 0.02 m and 0.035 m according to the species of fish in the tank. The fish were encouraged to move around using a bright handheld LED lamp array designed for video lighting. The tank was placed on a concrete floor on a ground floor of the building; small vibrations such as camera motors, if transmitted into the tank via physical contact, can bias the result (Moisy 2009).

**Figure 1.**
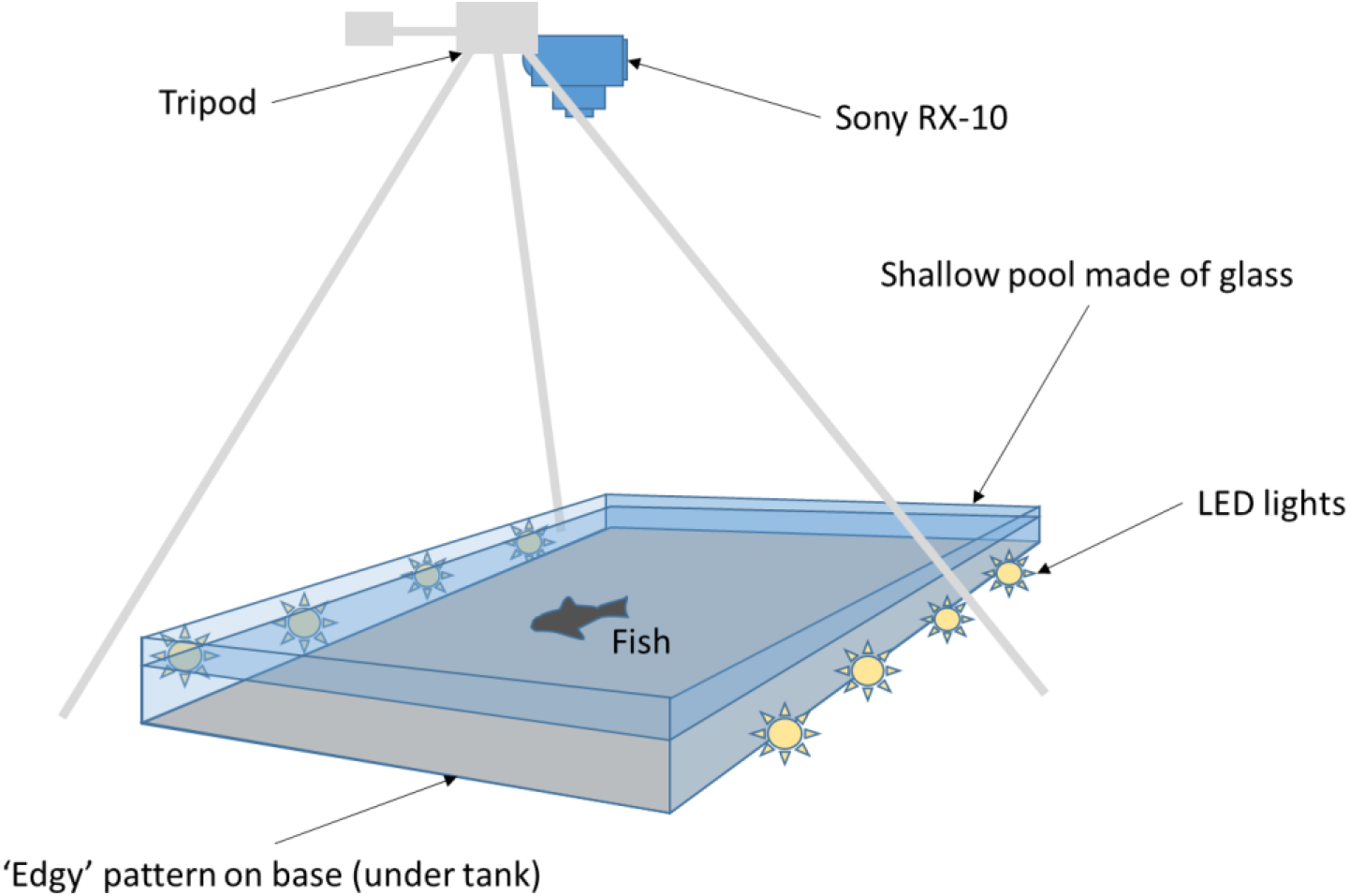
Schematic of basic apparatus. The camera is placed 1m above the tank surface on a tripod facing down onto the surface of the pool. The pool had a water depth of 0.02 m to 0.03 m. The tank was made using 6 mm standard float glass and silicon sealant often used in aquarium manufacture. The Sony RX-10ii camera that was used has a maximum video frame rate of 1000 fps, and so the lighting was strong and provided by standard aquarium (waterproof) LED lamps which do not vary (throb) with mains alternating power (at 50 hertz). The lights were placed below the water surface to avoid reflection from waves on the surface.

### Theoretical operation

Following Moisy et al (2009), the expected results of fish movement would be displayed on the camera as displacements in the basal pattern (Fig. 2). As a fish body moves in the water a wave is built up with components on both sides of the body, a pressure side (where the wave is positive above the mean surface level) and a suction side (where the wave is negative in depth with respect to the mean surface of the pool). The waves all move in the same direction as the fish body. Imaginary rays of light between the camera and the basal pattern are deflected by the air-water interface in predictable ways and the height of the water surface can be calculated (Moisy et al. 2009).

**Figure 2.**
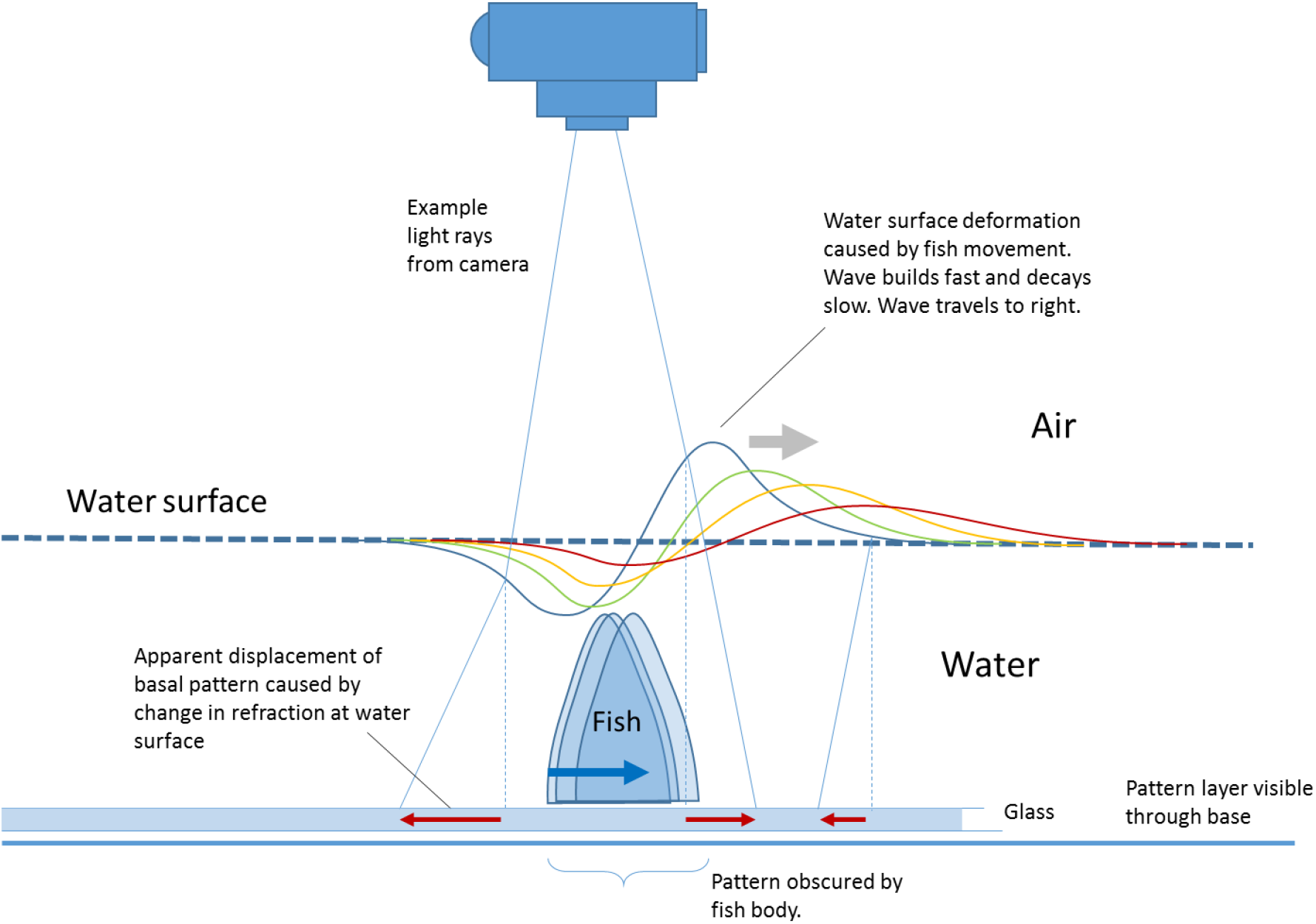
Theoretical operation of the schlieren method with a moving fish. The fish body is shown in cross-section and moves to the right in this diagram. The surface of the water is shown deflected into a wave of decreasing amplitude in blue, green, yellow and red curves as it decays in time. The wave group moves off to the left, in the same direction as the fish body. The wave is initially made by a high relative pressure caused by the force of the fish body trusting into the water on the left and a suction force as the body moves away and leaves a lower pressure region on the right of the body. Imaginary rays from a camera are shown as thin blue lines and show how the light from the basal pattern is deflected in various directions by the surface of the water as it is deflected in the waves. The diagram is not to scale and the waves would usually be lower amplitude than shown.

### Experimental subjects

The primary experimental subject was a male Chinese hill stream loach (Pseudogastromyzon myseri). The other species examined were a female butterfly loach (Sewellia lineolata), lizard fish (Homaloperoides tweedie) and a weather loach (Misgurnus anguillicaudatus). These are all related Cypriniform fish species in the families Cobitidae, Gastromyzontidae, and Balitoridae. They all inhabit South East Asia and China. The subjects had been acquired through the domestic aquarium trade, lived in our aquarium for six months, and continue to show no adverse effects several months after participation in the observations described here. We chose these fish to represent a range of swimming styles determined by morphology. The Sewellia are most elaborately adapted to life in fast flowing water with very large fan shaped pectoral and pelvic fins which overlap and form part of a body sucker system, whilst the weather loaches are the most conventionally fish shaped and can swim sinuously like an eel (as their species name suggests), but they can also swim by almost exclusively using the caudal fin (Thunniform gait). All these fish are thixophilic and very robust in terms of touching the substrate and squeezing under rocks and so forth, which ensured they were relatively comfortable with physical handling and swimming naturally in the shallow water of the observation tank.

### Existing software

We followed the methods described by Moisy et al (2009) with respect to the tank, camera and background pattern. We used Ncorr v1.2 software (Ncorr.com) for Digital Image Correlation (DIC). Ncorr is an open source 2D MATLAB (mathworks.com) product written by Justin Blaber and provided by the Georgia Institute of Technology. Following Moisy et al. (2009) we used intgrad2 written by John D’Errico and provided on Mathworks File Exchange (mathworks.com) to provide the conversion from 2D displacement fields to estimated relative surface height.

### Custom software

We wrote a custom graphical interface in MATLAB (mathworks.com) to display the displacement fields and original video frames together, and extended this framework to identify the fish in each frame. We used a frame rate of 250 or 500 frames per second for several seconds in each sample and so an automated scheme for positioning of the fish was required and gave a more objective and reproducible result than manual identification of the fish in each frame. This proved to be a challenge for several reasons: 1) the background was designed to provide a high density of contrast-based edges for the DIC software to lock-on to, which were also picked up by standard object identification tools which then identified many false positive objects, 2) the fish object changed shape at each frame which prevented a simple shape-fitting algorithm from working effectively, 3) the fish were patterned and contained multiple visible regions (body, head, fins, etc.), and 4) the fins could be transparent and therefore parts of the background pattern observed through parts of the fish. Our software used a 3, or more, stage process of identification of pixels that were fish as opposed to background. We used the inbuilt functions of MATLAB’s (Mathworks.com) Image Processing Toolbox such as ‘bwboundaries’ to identify polygons around regions of similar pixel weight in various channels. We also tried other standard features of image analysis such as superpixels which form part of the Image Processing Toolbox. Once the fish overall was identified through an HSV (Hue Saturation Value) channel (we found the H channel most effective for this operation), sub components could be identified through another channel (such as one of RGB Red Green Blue image channels) or different thresholds on the same channel. We also investigated RGB variance for pixels, and variance of other single channels (i.e. red of RGB) based on the eight nearest neighbour pixels, or 16 next nearest. The process was supervised machine identification in that the first frame was used as a training frame to set the thresholding of the separate stages and then these thresholds and stages were used for the rest of the clip, with manual oversight, to ensure the settings produced meaningful results throughout the clip. We made and used a simple model of an undulating fish to compare patterns derived from our fish identification software with what we would expect to see from classical theoretical treatments of the way fish swim. We developed an algorithm for estimation of the spine position from the body outline of a fish (as the central chord), this process involved identification of initial estimate for the centroid using ‘regionProps’; a built-in function of MATLAB (Mathworks.com), then drawing concentric circles centred at the centroid and calculating the mid arc point of circle sections contained within the body polygon, and iterating this procedure many times. After several 100 iterations the cloud of midpoints (with illogical points removed) was used to interpolate a smoothed spine along the body of the fish. The spine could be manually ‘pinned’ at the ends to account for tail positions that deviated sharply from the general body curve (Fig 3).

**Figure 3.**
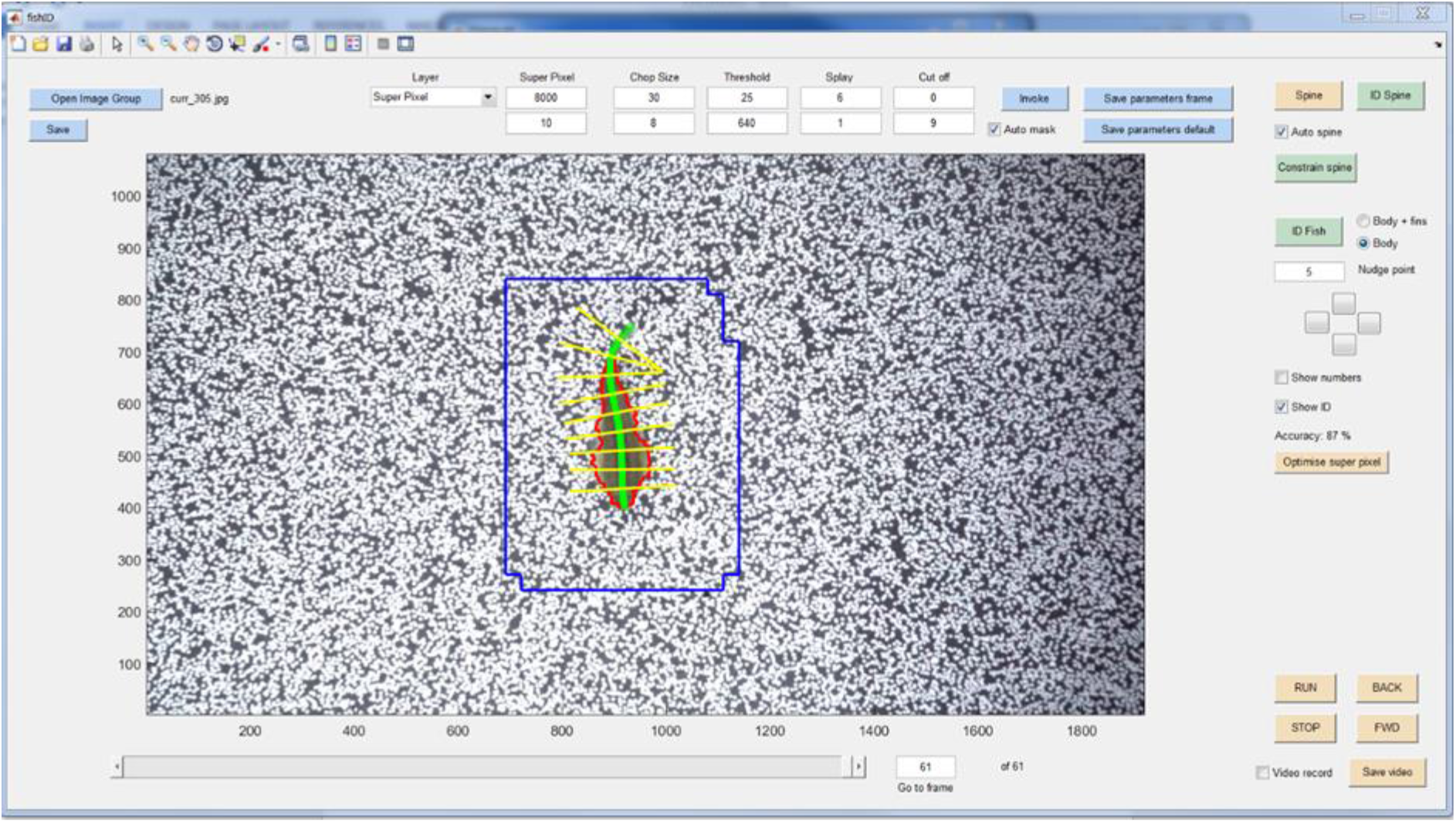
The custom built interface to identify the position of the fish body, the spine and the ribs, perpendicular to the spine. The blue box is the first pass automatic selection of the region of interest. The red polygon is the fish body outline, the green is the spine and the yellow are the ribs. The background is designed to be especially ‘edgy’ to allow the DIC (Digital Image Correlation) software to pick up small movements caused by deflection of the water surface. The fish is Pseudogastromyzon myseri. Axes labels are in pixel units.

At regular points along this spine, virtual vertebrae were defined (any number of vertebrae could be defined unrelated to actual vertebrae of the fish), and from these vertebrae ribs could be drawn that were perpendicular to the spine, and close to perpendicular to the skin of the body of the fish as they passed through it, and onward out into the areas beside the fish to measure the component of flow fields that were perpendicular to the side of the fish (Fig. 4).

**Figure 4.**
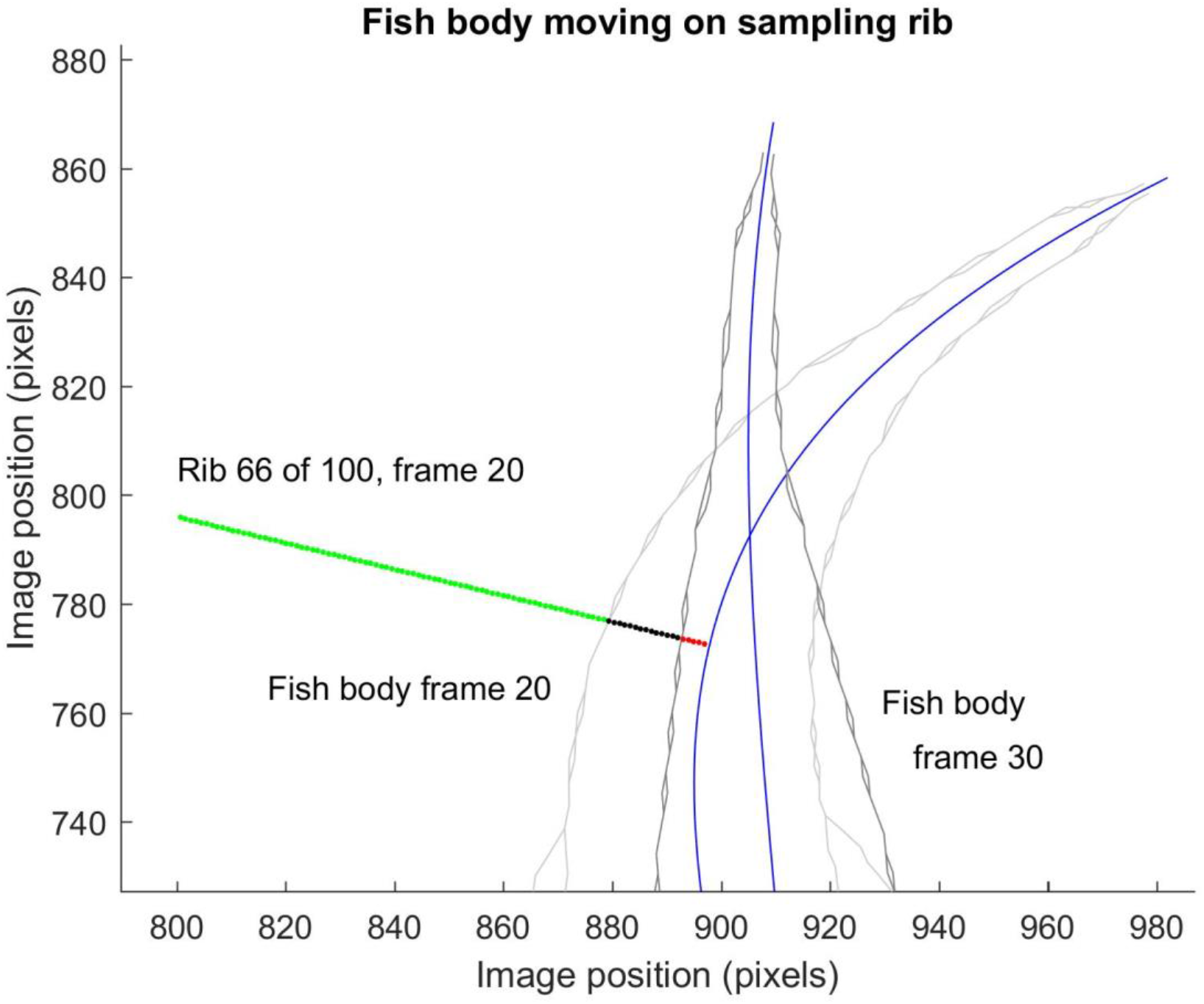
Illustration of the process of measuring the water surface deformation perpendicular to the skin of the fish. A rib in this case is shown as a green line, this is perpendicular to the spine at a positon which is 66% along its total length from the head to the tail. The black section is within the fish (and can be used to define the positon of the skin). As the frames progress the fish body moves off (light and then dark grey sets of lines – in this case drawn around body and body-with-fins which are convergent at the rear of the fish), but this rib is ‘left hanging’. By frame 30 the fish’s body has moved so that the rib intersects in only the red section. These hanging ribs can therefore show the development of movements that precede and follow each frame, including the lateral movement of the skin of the fish (albeit not the same points on the surface of the skin)..

## Results

The DIC software provided visualisation of the movements of the water surface inferred from the movements of the basal patterns (Fig 5). Two fish in the study produced good patterns, with two groups of results; hill stream loaches produced similar patterns, the lizard fish (Homalopteroides smithi) and the weather loach produced a different pattern. The false coloured overlaid images show that hill stream loaches (Pseudogastromyzon myseri) swim in this experimental set-up by a series of isolated thrusts, with waves produced by the body of the fish moving away from the fish at roughly 135 degrees with respect to the direction of swimming. Between these thrusts the body moved in a sinuous fashion consistent with the theoretical explanation of undulatory fish swimming movements. The weather loach, which is shaped more like a fish such as a mackerel (Scomber scombrus), when viewed in plan from the dorsal aspect, had an entirely different pattern of waves around its body during swimming (Fig 6). In this case all of the larger water waves were produced by the fish at its tail, as vortices in a turbulent wake. There were no waves evident along its body in a similar way to the hill stream loaches. The only wave not associated with the tail was a broad and feint bow wave which appeared to start behind the head of the fish and to travel at the same speed and direction as the fish, in the same way as the bow wave of a boat. Thus this swimming style, although superficially similar was revealed by the Schlieren method described here to be entirely different in terms of thrust pattern in the water and body location creating the main thrust.

**Figure 5.**
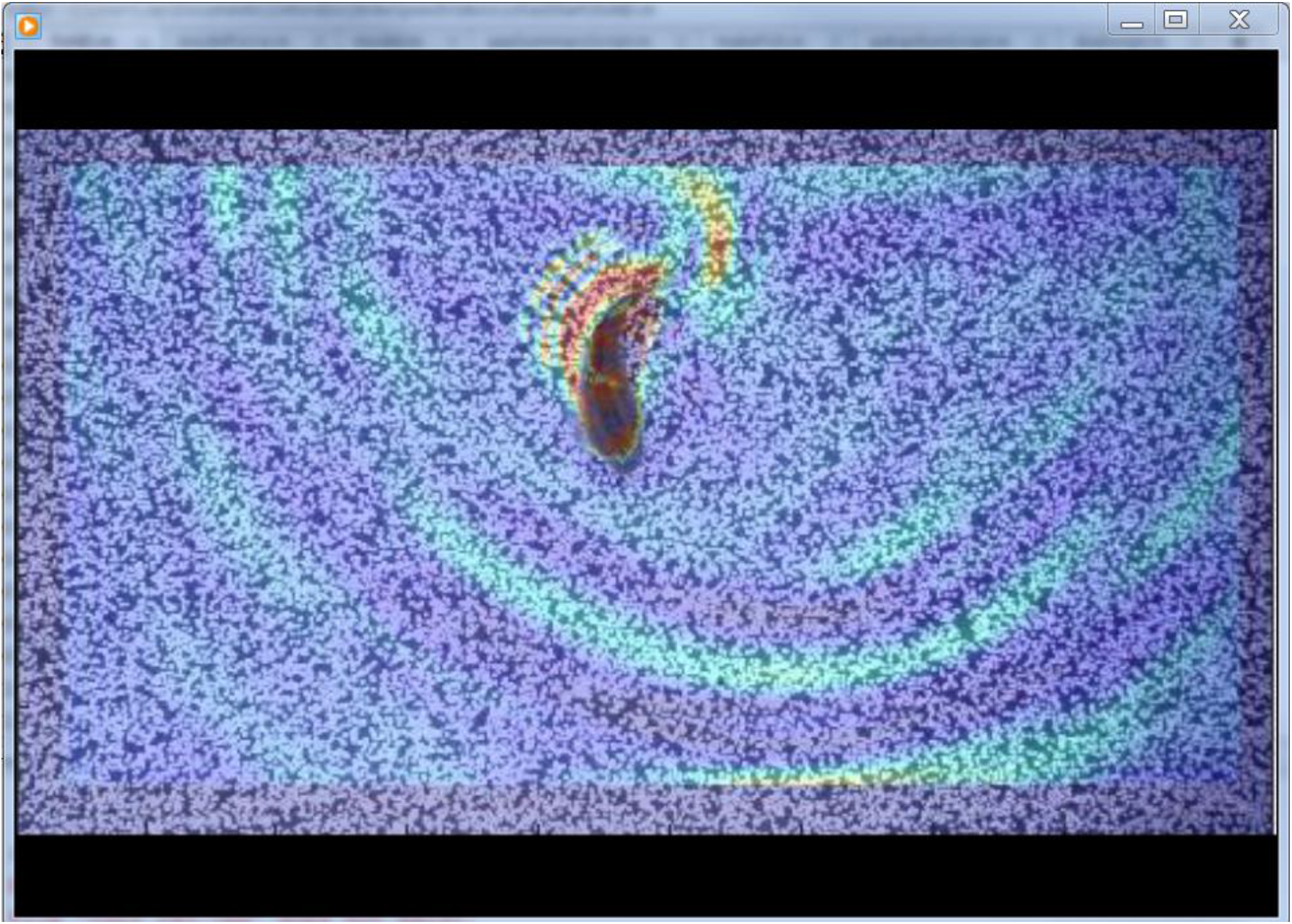
Video frame capture of the displacements of the basal pattern calculated using DIC (Digital Image Correlation) software of a swimming hill stream loach fish (Pseudogastromyzon myseri). The colours are the standard ‘jet’ colour map coloured with respect to velocity vector magnitude of the frame-to-frame displacements, with blue the slowest and red the fastest, and the alpha channel has been set to give a 50% transparency. The two red arcs to the left of the fish body are made by the two sides of a single positive pressure wave which is moving to the upper left. The arc to the right and above the fish is the residual effect of a previous stroke of the tail and caudal body area of the fish. The fish is swimming directly downward, and the frame rate was 250 fps.

**Figure 6.**
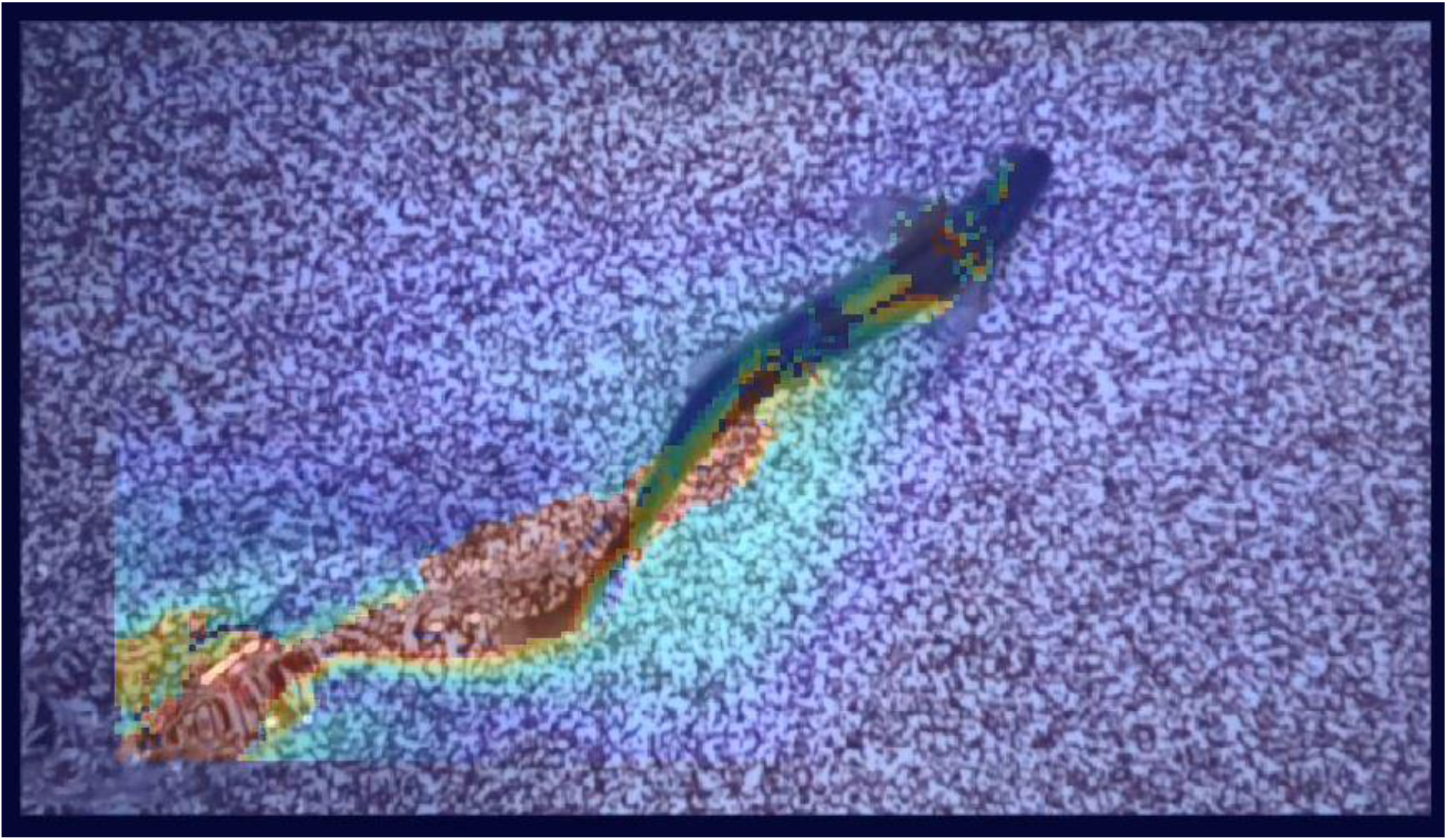
Video frame capture of the displacements of the basal pattern calculated using DIC (Digital Image Correlation) software of a swimming weather loach (Misgurnus anguillicaudatus). The colours are the standard ‘jet’ colour map coloured with respect to velocity vector magnitude of the frame-to-frame displacements, with blue the slowest and red the fastest, and the alpha channel has been set to give a 50% transparency. The fish is swimming toward the upper right of the frame, and the frame rate was 250 fps. Red areas at bottom left are the residual vortices in the turbulent wake of the fish. The fish body is in a sinuous form typical of an undulatory swimming gait with a wave of increasing amplitude moving along its body from head to tail. The red areas near the fishes body are caused by the movement of the fish body away from areas of basal pattern which it previously covered (they are not waves or indications of water surface changes but the DIC system picking up on movement of the fish itself). The bow wave is the feint light blue chevron shaped pattern roughly in line with the pectoral fins of the fish in this frame and emanating from around the point behind the fishes head between the junction of the pectoral fins and the main body.

The surface height reconstructed along the ribs of the hill stream loach shows the pattern of pressure and suction waves caused by the thrusting stroke of the fish. The wave can be viewed at an instant in time (a single frame) across a number of ribs, or as a function of time (multiple frames) on a single rib (Fig. 7 and Fig. 8). The wave can be seen to be in-phase along the body of the fish in a single instant and can be seen leaving the fish body before reaching the tail, which is contrary to the normal understanding of the thrust produced by the body and tail of an undulatory fish (Muller et al. 2002).

**Figure 7.**
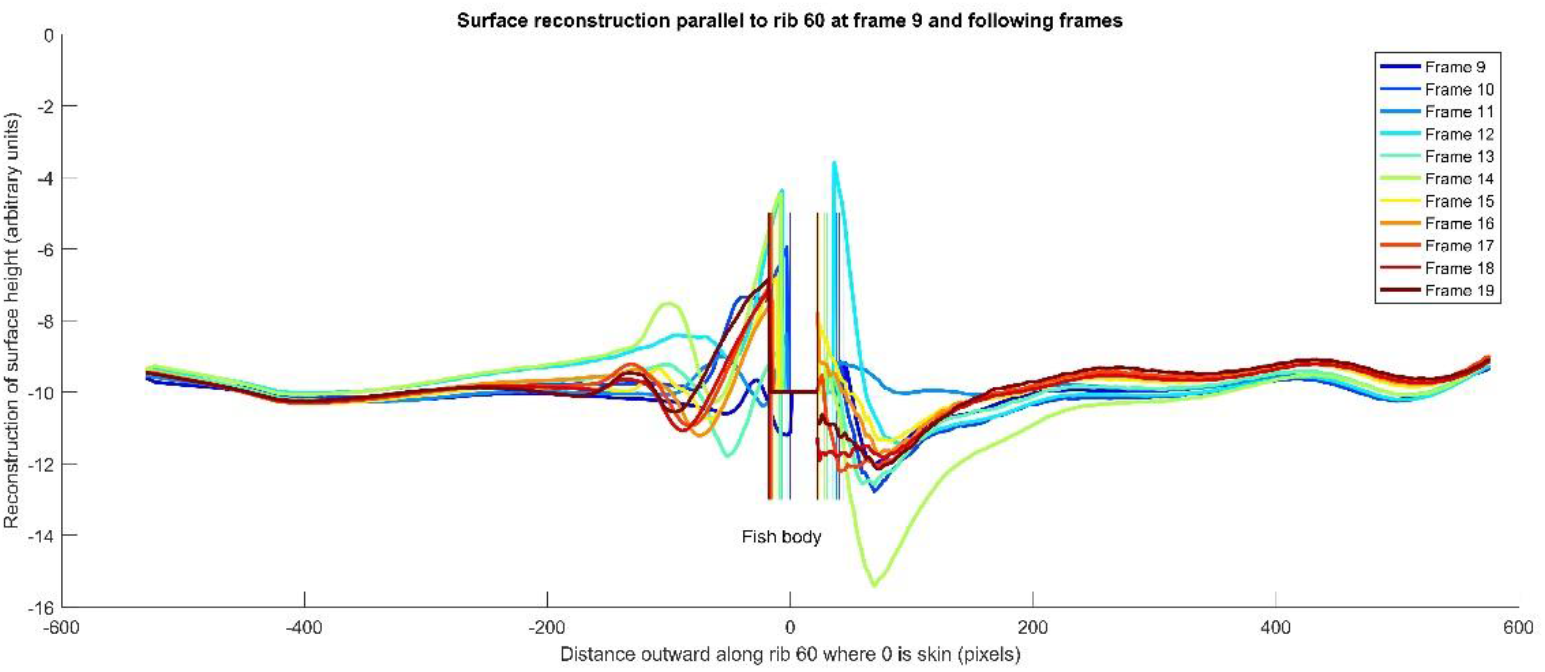
Water surface reconstruction along a single rib (rib 60 is 60% along the body of the fish from head to tail) during 11 frames of video (250 fps), x-axis scale is pixels, and y axis scale is arbitrary height units. During the frames the fish body moves to the left in this figure, as indicated by the vertical coloured lines which show the estimated position of the fish skin at each frame coloured to correspond with the frame as outlined in the key. The water surface can be seen to rise in a pressure wave on the left of the fish, and lower in a suction wave on the right. The waves are out of phase as they travel leftwards. The values very close to the fish (10-20 pix away) are unreliable as they are influenced by the movement of the body of the fish which was also tracked by the method described here. The fish was a hill stream loach (Pseudogastromyzon myseri).

**Figure 8.**
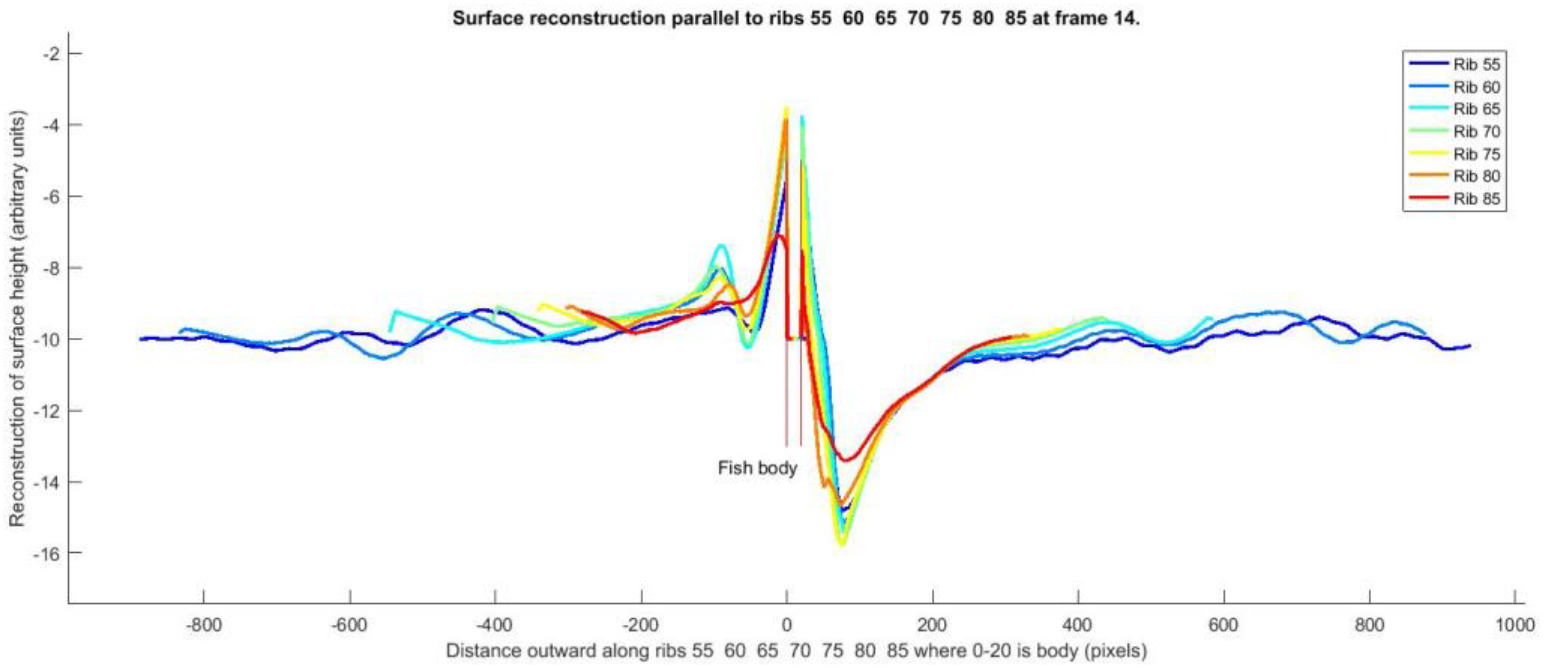
Water surface reconstruction along a 7 ribs at a single frame relative to the body edge of the fish (rib 60 for instance is 60% along the body of the fish from head to tail) x-axis scale is pixels, and y axis scale is arbitrary height units. The fish body is fixed during this frame and the estimated edges are shown by the red vertical lines in the centre. The fishes body was thrusting to the left in this figure (see Fig. 7). The water surface can be seen to rise in a pressure wave on the left of the fish, and lower in a suction wave on the right. The waves are in phase as they travel leftwards. The values very close to the fish (10-20 pix away) are unreliable as they are influenced by the movement of the body of the fish which was also tracked by the method described here. The fish was a hill stream loach (Pseudogastromyzon myseri).

The process of identification of the fish to link the DIC displacement fields with the position and orientation and movement of the fish and virtual ribs provided a detailed record of the fish body position and motion. For instance the interpolated position of the spine at each frame show the pattern commonly described as undulatory swimming gait (Fig 9).

**Figure 9.**
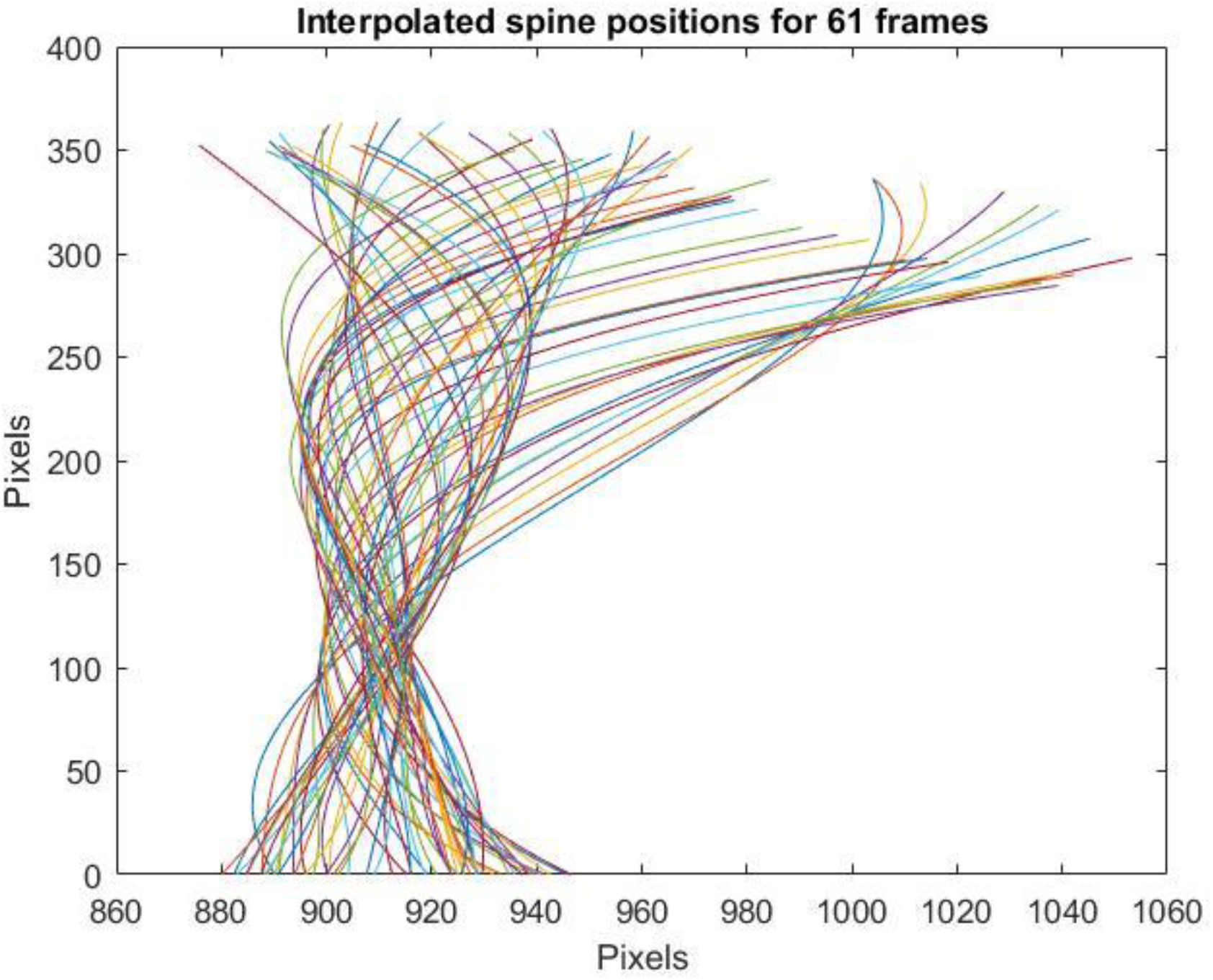
The interpolated spine positions derived from each frame of a 61 frame video sequence of the swimming fish. The axes are in pixels but are on a different scale to illustrate the detail of the movement by exaggeration of the lateral displacement. The fish is swimming directly downward in this case and all the spine positions at each frame are aligned in the vertical axis at the nose. The fish was a hill stream loach (Pseudogastromyzon myseri). The group of lines which move out to the right of the figure are the exaggerated movement of the C-start when the fish began swimming. A key feature is the point at around 100 pixel on the y-axis where all the spine positions cross over within a small area (the smallest area of convergence through time). We call this the hydrodynamic pivot, where the lateral movement of the fish was at a minimum and the movement in line with the swimming motion could therefore be most accurately estimated.

## Discussion

In comparison to alternative methods of visualisation of the wake of swimming fish the Schlieren method outlined here has some major advantages. There are also limitations and disadvantages. The major advantage is that is it inexpensive, simple and quick to set up when compared with systems such as laser PIV (particle Image velocimetry) (e.g. Druker and Lauder 1999). While Laser based PIV can give three dimensional flow patterns of exquisite quality, the lasers can be damaging to the eyes of fish, particles need to be introduced into the water, the images are usually restricted to the wake well after it has left the fish, it is only used away from the surface and substrate, and the experimental apparatus is restrictive with respect to the volume of water being examined. These limitations mean the experimental design is often far from one with which the fish is familiar and its behaviour may be impacted. Other methods that have been used such as introduction of dye into the mouth of fish, and dye streams in the water, while being cost effective and simple to use are potentially stressful to the fish, or in the case of dye streams in the water very difficult to apply consistently as required and the dye can obscure important parts of the fish body (e.g. Aleyev 2012). For fish which are very familiar with swimming in shallow water, close to the substrate, the Schlieren method here is a reasonable first choice. The fish swim in water with no additives, near to surface and substrate in lighting that is strong but well within their normal daylight tolerance, and the wake is visible close to the point of origin on the body of the fish. The primary limitation is that this is a 2D system and the body of the fish can obscure useful information as it moves to cover the background image. This is not so much of a problem for the identification of displacement fields and for the qualitative visualisation of waves, but it does cause the integration of the displacement field to be inaccurate as the constants of integration become corrupted by the fish as a displaced field. We generally found that the constants of integration were not consistent (Figs 7 and 8) We focused on the displacement fields as indicative of relative thrust as these were directly calculated by the DIC software and were consistent and accurate, when looking at the water surface deflection pattern, we focused on the relative surface height rather than the absolute height, which was sufficient for demonstrating the differences in phase that we required. Some calibration would be required if the apparatus was used to calculate absolute surface heights, or forces.

The swimming style we observed is different to styles that have otherwise been mentioned in the scientific literature, to our knowledge. The literature on fish swimming is extensive. Webb (1994) provided a wide classification of undulatory gaits of fish (based originally on the types introduced by Breder (1926)) and explained the balance of lift and drag experienced by many benthic and bentho-pelagic species, but only suggested that benthic fish suppressed their sustainable swimming gaits in order to either hold on to the substrate or use their fins (especially pectoral fins) to create downforce in order to maintain station in high flow. Small rheophilic fish often have teardrop shaped bodies as opposed to larger fish and mammals which are aerodynamically shaped (cigar shaped) identified as a potential parallel evolutionary trait associated with life in high speed flows (Hora 1952). This is likely to increase drag for a given body volume but lift is likely to be a more challenging force than drag (Vogel 1994) while holding station (or moving forward while remaining in contact with the bed). The forces on conical bodies in currents (Denny 2000) suggest that the teardrop shape of the rheophilic fish may be a compromise between ensuring the centre of lift and the area of suction, or other attachment, are aligned sufficiently to be effective. A teardrop shape includes a large relative head cross-section, by definition, and body flexibility is necessarily reduced in comparison to the more elongate fish. Small rheophilic fish may be constrained to use a short powerful in-phase pulse mode of swimming because the body length to body width ratio is insufficient to allow for an efficient undulatory swimming stoke.

One aim of the study described here was to investigate if it was possible to detect the likely thrust pattern of a fish from very precise photography alone. In this case the study here could be used as a type of calibration or verification of such a technique. With high speed cameras of high definition in normal lighting and with image processing software to automatically and reliably position the body of the fish it may be possible to infer thrust pattern and value. To this end we developed a dynamic model of a swimming fish based on the classical concept of the vector decomposition of a moving plate through water (Lighthill 1971). We compared this model with the results from this study. For instance we could assess the likely impact of a large immobile head on the swimming gait of a model fish. The model proved a valuable aid to understanding, and reliable explanation of the impact of an immobile head for instance. However, it exposed a limitation of the techniques in this study; that is the accuracy with which it is possible to measure the position of the body of the fish. Since the fish is in shallow water, and we are dependent on deflection of the water surface to produce the patterns of the substrate to show thrust, we are also victim of the same distortion in our attempts to measure the exact position of the fish. We found that the movement of the fish through the water appeared very similar to the expected sinuous gait of an undulatory fish, but the thrust patterns showed clearly that it made slight accelerations at key points during the stroke to produce pulsed thrusts. We were unable to measure the position of the body accurately enough to reproduce this pattern in our models due to distortions of the water surface near to the body. Future work with a submerged fish and submerged camera may prove effective in quantifying this pattern of movement. One unexpected benefit related to this was evident; the Sewellia species of butterfly loach have complex body colour patterns. The Sewellia lineolate has a body pattern very similar in many ways to the edgy pattern of the artificial background used in this study. This meant that the DIC software picked up the precise movements of the body of the fish, including deformations such as one would expect in a stress pattern example. This has potential for identifying precisely the locations on the body of the fish which are deformed in the swimming stroke, in effect measuring the skin of the fish as a stress deformed solid. Again this technique would be improved by using a submerged camera and a fish in slightly deeper water as the water surface deformations were noise. While outside the scope of this present paper these techniques might prove effective in many cases of analysis of swimming fish.

